# 3D Mapping Reveals Network-specific Amyloid Progression and Subcortical Susceptibility

**DOI:** 10.1101/116244

**Authors:** RG Canter, H Choi, J Wang, LA Watson, CG Yao, F Abdurrob, SM Bousleiman, I Delalle, K Chung, L-H Tsai

## Abstract

Alzheimer’s disease is a progressive, neurodegenerative condition for which there is no cure. Prominent hypotheses posit that accumulation of beta-amyloid (Aβ) peptides drives the neurodegeneration that underlies memory loss, however the spatial origins of the lesions remain elusive. Using SWITCH, we created a spatiotemporal map of Aβ deposition in a mouse model of amyloidosis. We report that structures connected by the fornix show primary susceptibility to Aβ accumulation and demonstrate that aggregates develop in increasingly complex networks with age. Notably, the densest early Aβ aggregates occur in the mammillary body coincident with electrophysiological alterations. In later stages, the fornix itself also develops overt Aβ burden. Finally, we confirm Aβ in the mammillary body of postmortem patient specimens. Together, our data suggest that subcortical memory structures are particularly vulnerable to Aβ deposition and that functional alterations within and physical propagation from these regions may underlie the affliction of increasingly complex networks.

**Author Contributions:** RGC, KC, L-HT, ID conceived of the work and planned the experiments.

RGC, HC, JW, LAW, CGY, FA, SMB performed experiments and analyzed data.

HC built the custom microscope.

RGC, L-HT, KC, ID wrote the manuscript.

## Introduction

Cognitive impairments attributable to Alzheimer’s disease (AD) will affect millions of individuals in the next decade, however the etiology of the disease remains unknown.^1^ The results from decades of research support the hypothesis that the accumulation of toxic amyloid-beta peptides (Aβ) in the brain contributes to the onset and progression of AD.^2,3^ The hypothesis was initially based on the discovery that mutations in the Aβ precursor protein (APP) and its processing enzymes cause autosomal dominant, inherited, familial AD (FAD).^4^ Subsequent preclinical studies demonstrating that the Aβ peptides resulting from these mutations induces synaptic loss and neuronal death *in vitro* and *in vivo* have further established the acute toxicity of aggregate forms of the peptide on neural substrate, which suggests they directly influence the neurodegeneration observed in different subtypes of AD.^5–7^ In addition to cellular harm, amyloidosis in murine models of AD contributes to AD-like memory impairments, hippocampal synaptic loss, and electrophysiological alterations -- changes that are also observed in human patients.^8–10^ This data not only links lesions to dementia in FAD, but also provides a basis for understanding the etiology of the sporadic form of the disease. Despite the link between amyloid toxicity and neurodegeneration, the precipitating events that trigger Aβ accumulation, deposition, and propagation which initiate AD remain unclear.

One impediment to understanding Aβ deposition has been identifying the brain regions most vulnerable to Aβ plaques. The first studies using postmortem brain sections from AD patients’ brains suggested that primary accumulation occurs in the neocortex with subsequent spread of aggregates to deeper structures implicated in learning and memory.^11^ However, due to the use of post-mortem specimens, these reports could not definitively link evidence of amyloid deposition to the eventual development of AD. Although recent advances in positron emission tomography (PET) imaging have enabled longitudinal patient imaging studies that confirm the importance of cortical Aβ in the diagnosis and prediction of AD progression,^12,13^ they have largely not reported the contributions of deeper structures to Aβ load and spread.^13,14^ Intriguingly, while cortical Aβ does correlate with disease status in patients,^15,16^ many cognitively healthy individuals also have high levels of cortical Aβ.^17^ The discrepancy between cortical accumulation and cognitive impairment suggests that unexplored Aβ-induced changes in the brain contribute to memory-related alterations in AD.^18^

Increasing evidence suggests that distributed memory networks are particularly affected by Aβ in AD. Some lines of investigation demonstrate alterations in high-level cortical networks like the default mode network (DMN),^14,19^ while other recent studies show core alterations in fundamental deep memory structures that are part of the limbic system.^10,20–22^ Though the cognitive and memory networks implicated by patient imaging somewhat overlap, there are distinct brain areas and functions that the networks do not share, leaving the hypotheses at odds. Whether the divergence arises from technical considerations or has unaddressed biological significance remains unknown, and still unexplored is the question of whether specific areas show differential susceptibility that is not seen with bulk detection. We hypothesized that some of the discrepancies in observed network susceptibility may be due to the difficulties in staging patient AD progression, which can be affected by variable pathological load,^18^ as well as by socioeconomic, lifestyle, and genetic factors.^23^

To overcome the variability in human disease, in this work, we capitalize on the temporal precision of murine models harboring FAD mutations to map amyloid progression at high-resolution throughout the brain.^24,25^ We use optimized techniques for whole-brain SWITCH immunolabeling^26^ to create an unbiased, spatiotemporal map of Aβ deposition. Our observations reveal early subcortical susceptibility to amyloidosis in regions connected by the fornix, such as the mammillary body, which shows concurrent functional alterations alongside aggregate formation. The fur-dimensional (4D) nature of the data also reveal potential mechanisms of propagation along white matter tracts and suggests that pathological changes within long-range projections that connect memory and cognitive networks may contribute to disease progression.

## Results

Using optimized system-wide control of interaction time and kinetics of chemicals (SWITCH) techniques (Figure 1a) that enable homogenous, whole-brain immunolabeling,^26^ we created a spatially unbiased, temporally precise map of amyloidosis in a murine model of AD. We chose the 5XFAD transgenic mouse line because it is a widely-used model of AD-like pathology that shows an age-dependent increase in Aβ aggregates and alterations in hippocampal physiology.^8,25^ Although the SWITCH techniques are widely applicable to many antibodies and target proteins,^26^ we discovered that we needed to further refine the buffer systems to achieve homogenous whole brain labeling with our chosen amyloid antibody. In our optimized SWITCH system, we switch pH and ionic strength of the buffers to actively modulate antibody-antigen binding kinetics^27^ In the first step, we use high pH and high ionic strength buffer to slow the binding reaction so that antibodies can penetrate deep into the sample. Next, the buffer is titrated to a physiological pH and ionic strength to allow antibodies to rapidly bind to targets. By employing this system, we were able to achieve homogenous labeling throughout thick tissue sections (Supplemental Figure 1). To further optimize the conditions for our chosen amyloid antibody, we used aged animals with the expectation that they would show wide-spread aggregate accumulation based on previous reports.^25,28^ Using the optimized SWITCH protocols to label a whole brain, we observed homogenous immunolabeling of Aβ aggregates throughout the entire tissue in aged mice (Supplemental Figure 2, Video 1). These large tissue volume techniques allow analyses of intact specimens without the loss of critical information to the size limitations and directional segmentation that are inherent in sectioning.^29^ To our surprise, careful examination of the entire brain revealed that, in addition to deposition in cortical and hippocampal regions known to be affected in AD, significant accumulation of Aβ was also evident in subcortical regions.

**Figure 1:**
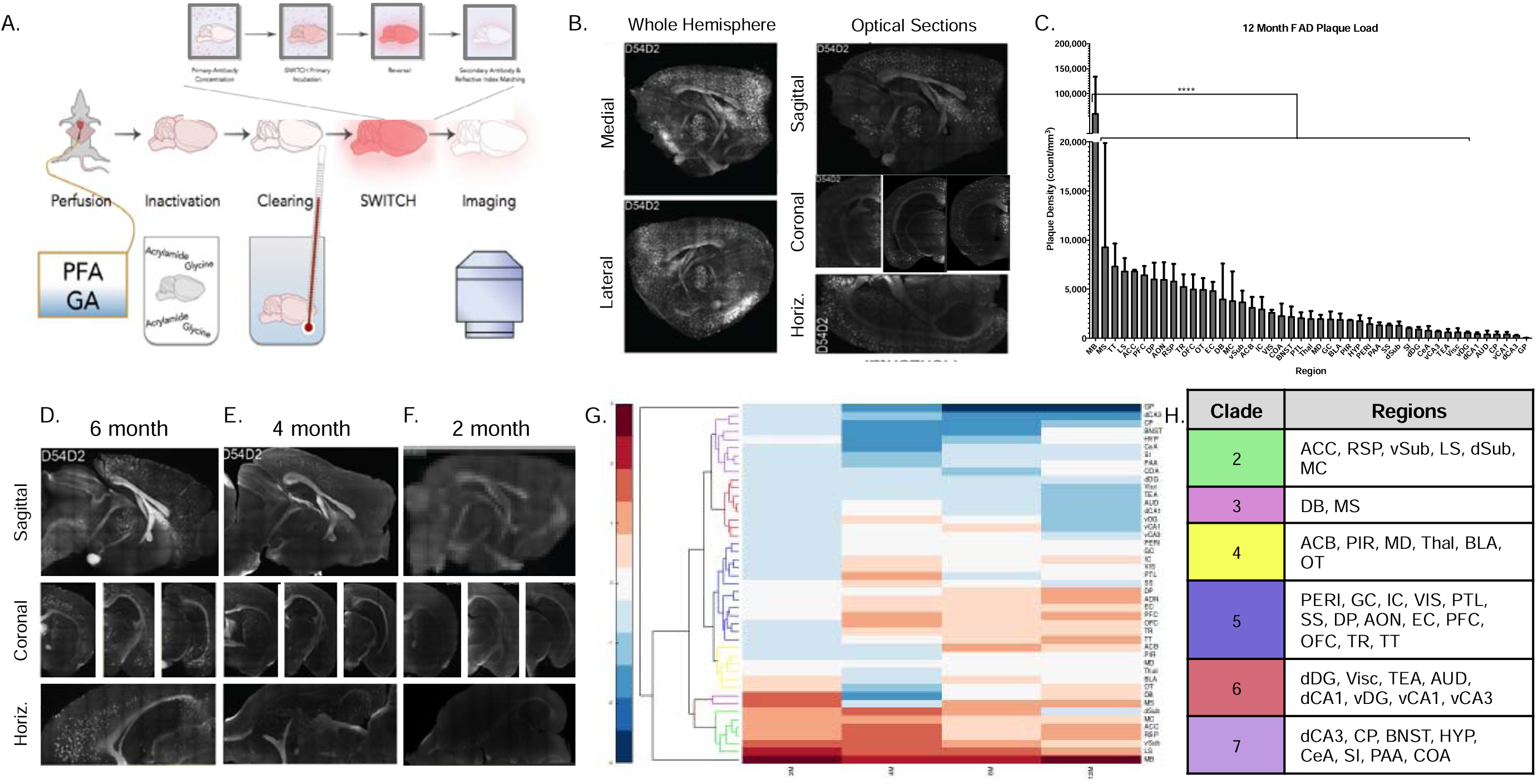
SWITCH labeling reveals network propagation of amyloid pathology. A) Diagram of protocol for our optimized SWITCH labeling. B) Representative images of the 12 month 5XFAD brains. C) Amyloid deposit density rank-ordered by region in 12 month aged 5XFAD mice. D-F) Representative images of amyloid labeling from D) 6 month, E) 4 month, and F) 2 month 5XFAD mice. G) Clustergram of the hierarchical clustering by density across the four time points examined. Coloring of the dendrogram represents the fifth level from the top and these clades contain regions with similar amyloid deposition patterns. H) Lists of regions contained in colored groups from panel G. *p ≤ 0.05; **p ≤ 0.01; ****p≤ 0.0001; Graphs report Mean ± Standard Deviation unless otherwise noted in the legend text. *Abbreviations: ACC, anterior cingulate cortex; RSP, retrosplenial cortex; vSub, ventral subiculu; LS, lateral septum; dSub, dorsal subiculum; MC, motor cortex; DB, diagonal band; MS, medial septum; ACB, nucleus accumbens; PIR, piriform cortex; MD, midbrain; Thal, thalamus; BLA, basolateral amygdala; OT, olfactory tubercle; PERI, perirhinal and ectorhinal corticies; GC, gustatory cortex; IC, insular cortex; VIS, visual cortex; PTL, posterial parietal association areas; SS, somatosensory cortex; DP, dorsal peduncular area; AON, anterior olfactory nucleus; EC, entorhinal cortex; PFC, prefrontal cortex; OFC, orbitofrontal cortex; TR, postpiriform transition area; TT, taenia tecta; dDG, dorsal dentate gyrus; Visc, visceral cortex; TEA, temporal association areas; AUD, auditory cortex; dCA1, dorsal CA1, vDG, ventral dentate gyrus; vCA1, ventral CA1; vCA3, ventral CA3; dCA3, dorsal CA3; CP, caudoputamen; BNST, bed nucleus of the stria terminalis; HYP, hypothalamus; CeA, centromedial amygdalar nuclei; SI, substantia innominata; PAA, piriform amygdalar area; COA, cortical amygdalar area*.

To determine the reproducibility of the protocols and replicate the unexpected labeling, we repeated the procedure in a cohort of 5XFAD animals aged 12 month (12M) (Figure 1B, Video 2d). In the 12M animals, we observed substantial plaque burden throughout the brain. Region specific quantification from white matter-based hand annotation of three-dimensional (3D) images (Supplemental Figure 3) demonstrated area-specific accumulation in AD-associated areas (Figure 1C). The regions harboring Aβ included the retrosplenial cortex (RSP), hippocampus (HPC), subiculum (SUB), and pre-frontal cortex (PFC), and importantly, aggregates were largely absent from areas thought to be relatively spared from amyloidosis in AD like the caudoputamen (CP) and midbrain (MD) (D’Agostino and Pearson normality test; K2 = 94.61, p < 0.0001, passed normality = No; Friedman test; Q = 114.2, p < 0.0001). This suggests that our labeling techniques are able to homogenously and specifically label Aβ aggregates, and that accumulation of the peptide is restricted within particular brain areas. Unexpectedly, the highest aggregate density by rank in the 12M animals appeared within specific subcortical regions, namely the mammillary body (MB) and septum (SEPT), which confirmed our initial observations.

Although the MB and SEPT had the densest Aβ, with data from aged animals we could not determine whether this pattern emerged because aggregates in the 5XFAD animals develop uniformly throughout the brain over time, thus making these smaller structures appear more dense; or if these high-density regions show unique vulnerability to elevated levels of Aβ, leading to earlier or more significant aggregation. To observe how the pattern emerged, we used our optimized SWITCH protocol to analyze a time course of 5XFAD cohorts including mice aged for 6, 4, 2, and 1 months (6M, 4M, 2M, 1M respectively) (Video 2a-c, Figures 1D-G). Quantifying Aβ density in specific regions across multiple animals at each time point, we observed increasingly precise aggregation with decreasing age in a pattern that was consistent across individuals. At 6M, 5XFAD animals had region-specific aggregation, with areas harboring significantly different densities of amyloid. (Figures 1D, 1G; D’Agostino and Pearson normality test; K2 = 70.12, p < 0.0001, passed normality = No; Friedman test; Q = 68.56, p = 0.0171). To better understand the brain areas that contribute to region-specific aggregation in our data, we performed hierarchical clustering analysis, which revealed that at 6M, areas associated with the olfactory and cortico-limbic system (e.g. basolateral amygdalar complex [BLA], piriform cortex [PIR], entorhinal cortex [EC]) show most similar levels of Aβ (Supplementary Figure 4B; hierarchical clustering, single linkage-nearest distance). This is particularly interesting because it suggests that network-level distribution of Aβ is relatively uniform within functional networks and also identifies circuits that underlie behaviors known to show alterations in early AD.^30^

Looking at younger animals to determine the progression in the predictable, genetic murine models, we next saw that at 4M the data continues to demonstrate area-specific aggregation and our analyses show non-normal, non-linear pattern to the regional distribution (Figure 1E, 1G; D’Agostino and Pearson normality test; K2 = 51.67, p < 0.0001, passed normality = No; Non-linear Regression; y = −354.1*ln(x)+1200.9, R^2^ = 0.82465). The skewed data suggest that fewer regions have high burden, consistent with increasingly specific patterns of aggregation at the earliest disease stages. To determine what regions are affected at this 4M age, we looked at our clustering analyses (Supplementary Figure 4C), which revealed that regions homologous to the those in the DMN (e.g. restroplenial [RDP], anterior cingulate [ACC], and parietal cortices)^31^ are better correlated to each other, with the exclusion of olfactory and limbic areas from the cluster. To understand whether the DMN is the earliest affected network as suggested by many human functional studies,^32,33^ we next looked at 2M animals. At 2M, animals failed to demonstrate significant differences across brain areas (Figure 1F; D’Agostino and Pearson normality test; K2 = 105.2, p < 0.0001, passed normality = No; Friedman test; Q = 58.18, p = 0.1073), suggesting little to no specific aggregation patterns. However, our analyses revealed that while many areas did not cluster, there was correlated deposition in the 2M animals within a few areas (Supplementary Figure 4D). These key nodes of early aggregation were the MB, SEPT, and SUB – regions that connect the HPC to the rest of the Papez memory circuit.^34^ At 1M animals did not show discernable amyloid deposits (data not shown), suggesting that the deposition is not developmental and that the SUB, MB, and SEPT are uniquely susceptible to Aβ aggregation at the earliest stages of disease pathology.

One concern using transgenic models is whether the pattern is reflective of biologically interesting results, or a byproduct of the transgenic expression. To ascertain whether the subcortical aggregation pattern may be an artifact of the mouse transgene, we performed dual immunofluorescence for Aβ peptides and *in situ* hybridization for the transgenic mRNA to correlate the location of Aβ deposits with expression patterns of the transgene (Supplementary Figure 5A). Patterns of expression in 2 and 4-month animals looked similar and were combined, and the correlation analyses demonstrate that across the entire brain, transgenic RNA expression does not correlate with deposit location (Supplementary Figure 5B; ρ = −0.0728, p = 0.3588). The insignificance of this result remains when split by region (Supplementary Figure 5C). This data suggests that 5XFAD animals show biologically relevant, region-specific amyloid deposition. Combined with our clustering results (Figure 1H), our data show that the MB, SEPT, and SUB are particularly vulnerable to Aβ deposition. Over time, the aggregates appear in regions homologous to the DMN. Following the DMN, lesions are detected throughout the extended limbic system, before finally developing throughout the entire forebrain. Thus we propose that Aβ deposits start in susceptible subcortical structures, and spread to increasingly complex memory and cognitive networks with age.

The detection of aggregates alone is not sufficient to demonstrate that the Aβ we observe in the MB, SEPT, and SUB may contribute to Aβ-related cognitive decline because it is unknown whether subcortical Aβ, especially within diencephalic structures, induces the functional alterations thought to underlie early memory deficits in AD.^9,22,35,36^ In our data, the MB shows the densest Aβ at the earliest stages. The MB is downstream of both the SEPT and SUB,^37^ and previous reports suggest that it is related to recall memory in humans^38^ and has calcium-sensitive firing properties^39^ that may render it particularly vulnerable to Aβ-induced dysfunction. Thus, to test whether the observed aggregation leads to functional impairments within the subcortical structures, we performed *ex vivo* whole cell patch-clamp recordings in the MB in slices from 2M 5XFAD mice and littermate controls. 5XFAD MB neurons showed significantly increased excitability (Figures 2A-B, 2-way Repeated Measures ANOVA; Interaction: F_(10_,_450)_= 8.786, p < 0.0001; Genotype: F_(1,45)_= 9.902, p = 0.0029), which is consistent with previous reports of neuronal hyperexcitability in the hippocampus in response to Aβ^9,35^ Further examination showed that this excitability was not due to increased action potential threshold (Figure 2C, Unpaired Student’s t-test; t_(46_) = 1.146, p = 0.2577) or altered hyperpolarlization dynamics (Figure 2D, Unpaired Student’s t-test; t_(32)_ = 0.7107, p = 0.4824), but may be attributed to slightly increased action potential amplitude (Figure 2E, D’Agostino and Pearson Normality Test, FAD-: K2 = 11.09, p = 0.0039, passed normality = No; FAD+: K2 = 4.217, p =.1214, passed normality = Yes; Mann Whitney U; U = 87.5, p = 0.0497) and a reduced resting membrane potential (Figure 2F, Unpaired Student’s t-test; t_(40)_ = 4.807, p < 0.0001) in 5XFAD MB neurons compared to the WT controls. This is consistent with previous findings of altered resting membrane potential in non-pyramidal neurons of the HPC in a different murine model of AD,^40^ revealing further consistencies in amyloid-driven alterations, despite differing cell-types and circuit structures.^41^ Thus, our results suggest that early accumulation of A|3 the MB of the 5XFAD drives neuron-intrinsic hyperexcitability, which is characteristic of AD-related changes in neuronal activity and, in this early stage, may contribute to the onset of network dysfunction and memory impairment.^9,35,40^

**Figure 2:**
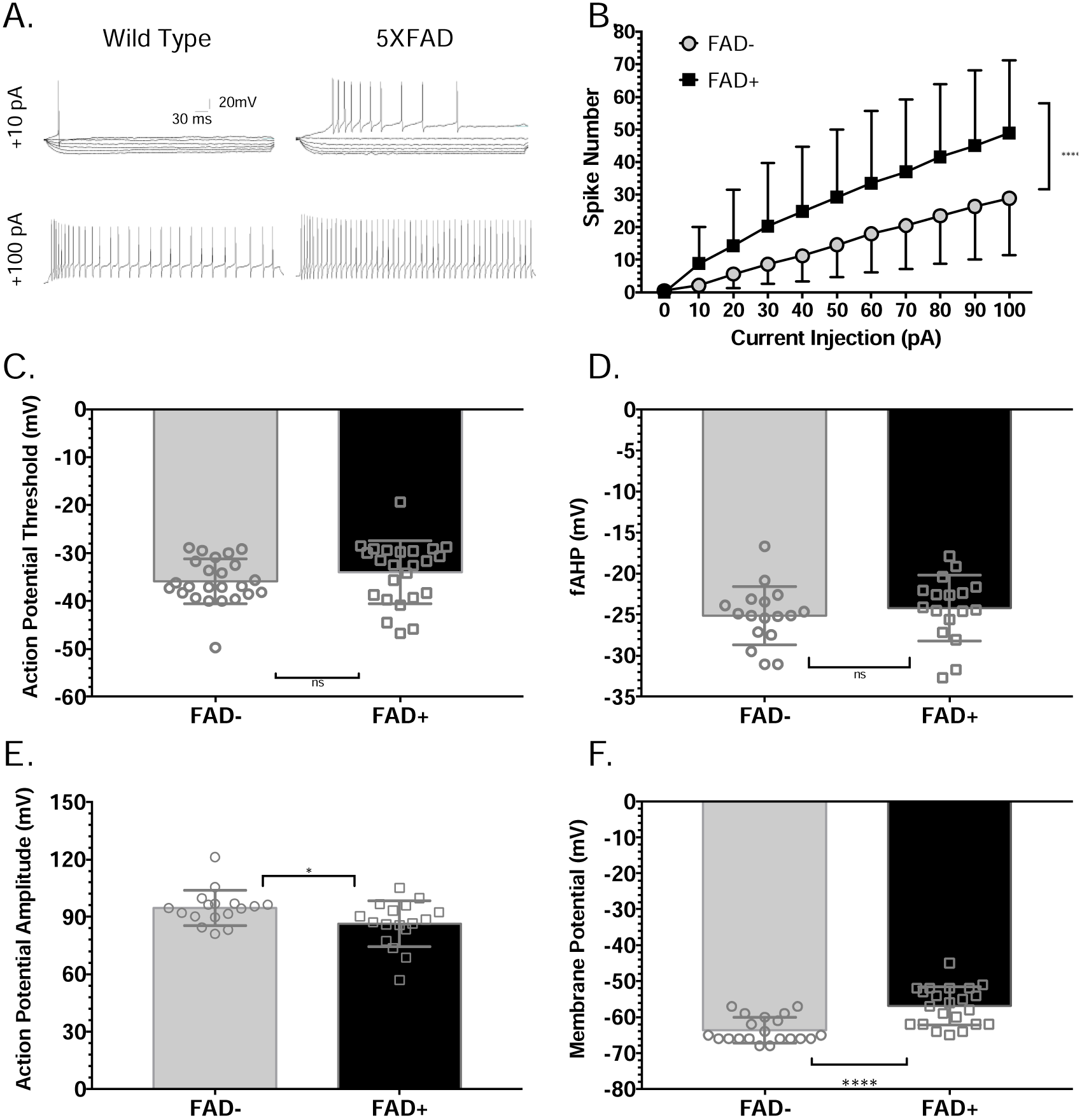
**The MB shows significant functional impairments** and loses its synaptic input. A) Representative action potential traces following current injection of 10pA (top row) and 100pA (bottom row) into MB neurons of 5XFAD animals (right column) compared to littermate controls (left column). B) Excitability curves showing increased firing rate of FAD+ across multiple steps of current injection. C) The action potential threshold of the cells does not change, neither does the D) after-hyperpolarization. E) The action potential amplitude is significantly decreased in 5XFAD animals and F) the resting membrane potential is also significantly decreased, both of which could underlie the excitability phenotype. Graphs represent Mean ± Standard Deviation.

Our results demonstrating that the MB is particularly susceptible to early Aβ deposition adds to the mounting evidence that implicates subcortical Papez circuitry in AD-related memory impairment in human patients,^21,42–44^ however the relevance of these findings is tempered by mixed reports about the involvement of MB in human mild cognitive impairment (MCI) and AD patients.^42,45,46^ Thus, to understand whether MB demonstrates Aβ lesions over the course of AD, we acquired a small cohort of post-mortem mammillary bodies embedded in paraffin blocks from the Netherlands Brain Bank. The blocks contained mammillary bodies from individuals that died at each Braak and Braak stage^47^ (Figure 3A). We cut traditional histology sections from the blocks (Figure 3B) that revealed numerous Aβ deposits in the MB of AD patients at each stage, with none present in the healthy individual (Figure 3C, 3D). This finding is consistent with older reports of amyloid pathology in the region.^43^ However, because the sections were thin and the deposits did not label strongly, we could not determine whether there was a stage-dependent increase in Aβ within the MB. To test this, we used SWITCH methods to clear and label the entire banked MB specimen from each individual. 3D reconstruction of the labeled specimens (Figure 3E, Video 3) revealed substantial Aβ burden in the MB of AD patients, but none in the MB from a healthy individual. Although the density of Aβ deposits did not correlate with NFT accumulation reflected in advancing Braak stage (Figure 3F), individual aggregate volume did (Figure 3G). This suggests substantial early Aβ deposition precedes overt dementia and that aggregates continue to expand as the largest ones in our small cohort were seen in the Braak V – AD patient. One additional interesting observation in the pathology sections was that there were numerous Aβ deposits in oligodendrocyte-rich regions of the MB (Figure 3H), suggesting a relationship between white matter tracts and Aβ deposition in this region.

**Figure 3:**
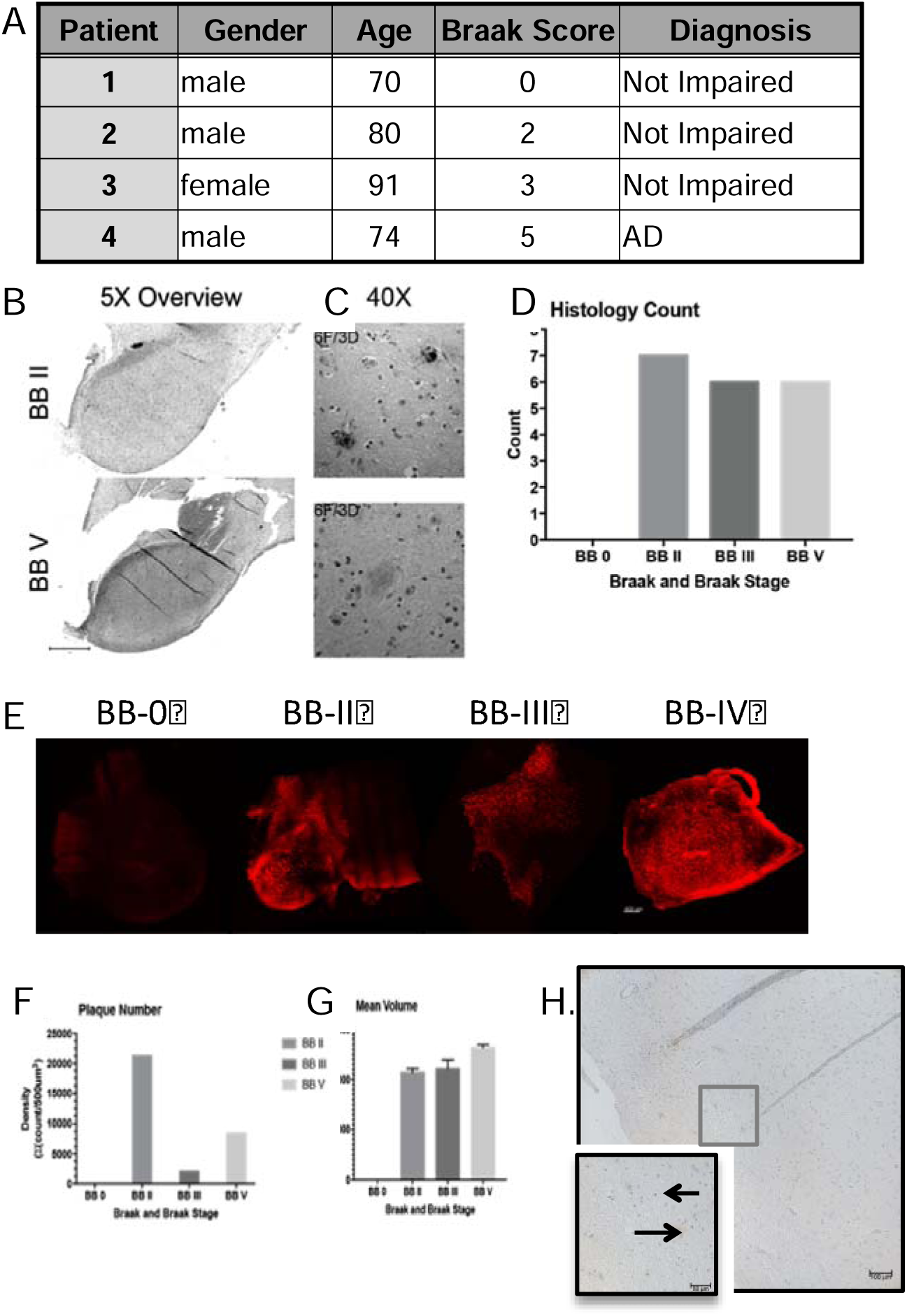
Human mammillary body and associated white matter show AD-related amyloid burden. A) Information on the patient samples used for the experiments in this figure. B) Representative images of 5X tile scans showing the overall MB structure and C) representative high-magnification image of the plaque types used for quantification (right). D) Quantification of histologically identified plaques from pathology slide sections. E) Representative images of the large-volume MB samples labeled with D54D2. F) The plaque count does not reflect the severity of the disease, however the G) mean plaque volumes increase with disease severity in this small cohort. H) White matter tracts identified anatomically and pathologically by presence of oligodendrocyte nuclei show amyloid burden too.

To check whether this is reflected in the 5XFAD model, we investigated the Aβ burden in major forebrain white matter tracts. We did not observe Aβ in the white matter of 2M animals, but did observe consistent deposits in the fornix of the 6M cohort (Figure 4A, Video 2c). The fornix, which connects the SUB to the SEPT and MB, is the major output structure of the hippocampal formation. To determine whether this burden was unique to the fornix, we took 100um coronal sections from 6M 5XFAD brains (Figure 4B). Surprisingly, white matter tracts connecting major limbic structures showed significantly more accumulation than tracts connecting other regions (Figure 4C, Friedman Test, Q = 40.29, p <.0001), suggesting that the limbic white matter is uniquely vulnerable. To determine whether these aggregates formed along the outside fasciculated bundles, we performed higher-magnification imaging of the 100um sections (Figure 4D, Video 4). Interestingly, the 3D reconstruction demonstrates amyloid build-up within the myelin sheath, along the axonal path (Figure 4D, Z-plane). This pattern was specific to the limbic structures like the fornix, fimbria, and cingulum bundle (Figure 4E), excluded white matter tracts like the corpus callosum (Figure 4E) and was not present in the wild-type littermate control animals (Figure 4F). This data is particularly intriguing because in the 6M FAD there is considerable Aβ burden in the cortical regions connected by the corpus callosum (Figure 1D). Together, it suggests that limbic white matter tracts are uniquely affected by axonal Aβ in AD, and may explain structural alterations in these regions in human MRI^21^ and the progression of connectivity impairments in individuals at risk or harboring significant brain Aβ.^48–50^

**Figure 4:**
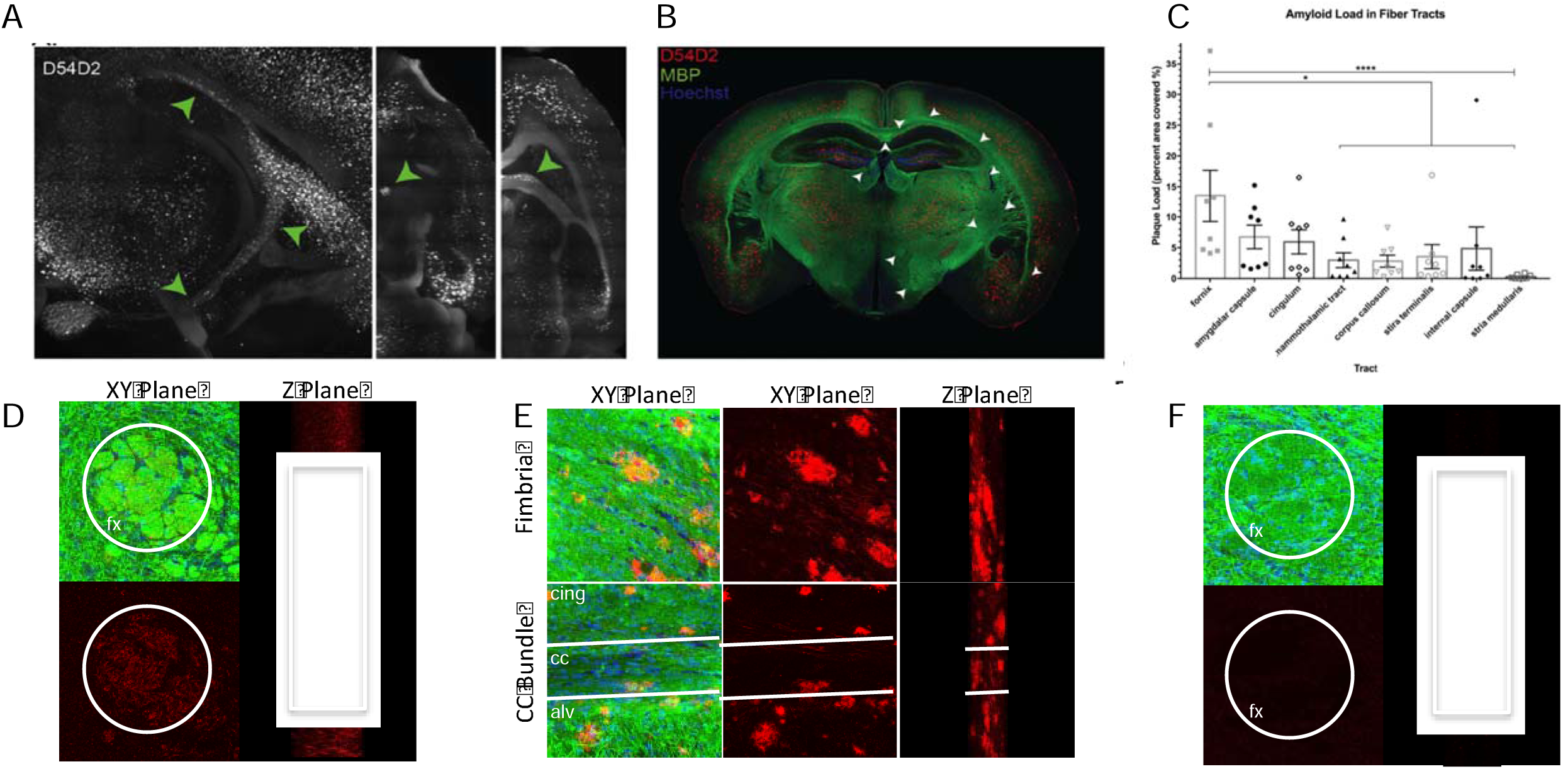
White matter amyloid deposits are specific to limbic tracts. Six month 5XFAD animals develop amyloid deposits within the white matter. A) Representative images of amyloid within the fornix visualized in the 3D rendered whole brain amyloid labeling in the six month 5XFAD cohort. B) Representative image of the white matter amyloid labeling in 100um sections from a second cohort of six month 5XFAD animals. C) Quantification of white matter deposits within several posterior white matter tracts. D) Representative images demonstrating labeling of amyloid within white matter bundles in the fornix. E) Representative images showing robust amyloid deposition within the fimbria, cingulum, and alveus, and less in the corpus callosum. F) There is no amyloid labeling within the fornix of non-transgenic 6 month aged littermates. *p ≤ 0.05; ****p≤ 0.0001; Green and white arrows point to white matter regions with observed amyloid. Graphs report Mean ± Standard Deviation. *Abbreviations: fx, fornix; cing, cingulum bundle; cc, corpus callosum; alv, alveus.*

## Discussion

Using optimized SWITCH whole-brain clearing and immunolabeling technologies, we have created a spatially unbiased map of the progression of Aβ deposits that revealed novel subcortical hubs of early-disease susceptibility in a mouse model of AD. Our findings demonstrate that areas connected by the fornix develop aggregates at the earliest phases of disease and that, of the identified vulnerable regions, the MB shows the most significant density of deposits across time. Although the MB is a critical part of the Papez long-term memory circuit connecting the hippocampus to the anterior thalamus,^34,51^ it has not been strongly implicated in AD.^34,42,52,53^ This is somewhat surprising because its major inputs, the subiculum and prefrontal cortex^54^, have been shown to demonstrate significant synaptic loss that correlates with memory performance.^55–58^ Furthermore, the MB is part of the head-direction system, a group of critical navigation regions including hippocampus, thalamus, and retrosplenial cortex that are affected in AD, and the pathology in which is thought to underlie spatial memory deficits seen in the disease.^59,60^ Integrating our findings into this framework, the present data indicate that the subcortical and diencephalic projecting components of the Papez circuit are particularly vulnerable in AD. Interestingly, some of the regions of the Papez circuit and head direction systems, like the septum, were initially implicated in early AD by the cholinergic hypothesis, which posited that loss of those neurons was causative in disease onset and progression^61^. Although the idea was overshadowed by the amyloid cascade hypothesis,^2^ our data, along with other recent evidence for Papez circuit dysfunction^50^ suggests that these subcortical structures and their white matter tracts are associated with amyloid pathology early in AD.^62^ In the present manuscript, we take this observation one step further to show that the onset of Aβ aggregation in the MB of the 5XFAD animals correlates with functional alterations. These data are consistent with network-level changes previously observed in the Papez circuit in AD patients and other amyloid models,^9,22,36,40^ and the significant findings at such an early pathological stage highlight the importance of continued investigation of the network alterations and functional changes in these structures at the initial disease phases. Because the regions in the basal forebrain are linked in our data, and other studies, to the hypothalamic diencephalic limbic structures, we suggest in-depth studies of these regions and their connections may lead to a better understanding of the mechanisms of dysfunction underlying AD onset and the ensuing progressive memory loss.

It has become increasingly important to understand the translatability of observations in animal models to human disease.^63,64^ Thus to determine whether the MB is also affected early in human AD, we used SWITCH techniques to confirm that Aβ aggregates are present in postmortem MB from Braak staged individuals with tau pathology and an AD patient. Our data investigating both traditional histology sections alongside large-volume labeled samples show significant Aβ burden that increases with stage, which is most evident using our novel technique. The data we obtained with traditional labeling methods was consistent with the literature^65^, however the large-volume images revealed that the histology sections did not show the full burden in the region. This observation highlights the power of these tools to enable a better understanding of pathological affliction through molecular interrogations of intact, precious samples. These observations of MB lesion and dysfunction in AD are of particular note because of the amnesia associated with other MB-related conditions. For example in Wernicke’s encephalopathy and Korsakoff’s syndrome, MB structural alterations were one of the first anatomical abnormalities correlated with worsened memory performance.^66–68^ More recent studies utilizing functional readouts and dietary interventions suggest that while the MB may be a primary site of sensitivity, thalamic, hippocampal, and cortical alterations also coincide with worsening memory.^67,69,70^ Interestingly, the interventions restore hippocampal connectivity to the diencephalic mammillary body and thalamus^69,70^, which are strikingly similar results to the experimental trials using deep brain stimulation of the fornix near the MB to enhance memory in AD patients.^71^ Together, this data suggest that dysfunction within the MB correlates with HPC abnormalities and memory impairment, like those seen in AD. Importantly, recent human studies correlating MB abnormalities to memory performance specifically link MB aberrations to impaired recall memory,^38,70^ the type of memory thought to be most affected in AD.^72,73^

In addition to a better understanding of the regions that show primary Aβ aggregation and promote the initial network destabilization, our data suggest mechanisms for Aβ propagation between connected brain regions.^74^ By demonstrating enhanced Aβ deposition within limbic white matter tracts, and unique axonal patterns of Aβ within the Papez and limbic circuit tract, we propose that specific disconnection of the core memory network underlies cognitive impairment in AD. This may be due to particular mechanisms of Aβ handling within primary neurons of the memory network, or the particular vulnerability of the cells to overproduction or mishandling of APP/ Aβ. Additionally, by showing that distinct networks are burdened by Aβ, with increasingly complex networks affected across age, we present a unifying view of early lesions and can provide a framework in which to better interpret human pathogenesis. Investigators have suggested that the DMN, limbic system, attentional systems, and brain stem may all be involved at the earliest stages of prodromal AD. The data presented in this manuscript suggest that these do not need to be disparate hypotheses, but instead represent different stages of network affliction that occur as the disease progresses. In the mouse model, 2M 5XFAD animals show core limbic affliction, followed by the DMN at 4M and typical limbic system at 6M. These observations suggest that early AD may be best staged by the network aberrations detected by functional MRI techniques. These alterations occur before overt memory loss,^33^ and with increasing evidence for restoration of network function as a successful treatment in AD models,^9,72,75^ our data may lay the ground work for network-progression staging to guide early-disease circuit interventions.

## Methods

### Animals

All mouse work was approved by the Committee for Animal Care of the Division of Comparative Medicine at the Massachusetts Institute of Technology. 5XFAD (Tg 6799) breeding pairs were acquired from the Mutant Mouse Resource and Research Center (MMRRC) (Stock No. 034848-JAX) and maintained as hemizygous on the BL6 background. Animals were group housed on a 12h light/dark cycle with Nestlet enrichment and sacrificed at the ages noted in the text.

### Mouse Tissue Fixation

Mice were deeply anesthetized with isoflurane (Isoflurane, USP, Piramal Healthcare, Andhra Pradesh, India) and underwent transcardial perfusion with ice cold 1X PBS (10X stock, Gibco, #70011-044) followed by ice-cold fixative made of 4% paraformaldehyde (32% stock, Electron Microscopy Sciences (EMS), Hatfield, PA, #15714) and 1% glutaraldehyde (10% stock, EMS #16110) in 1X PBS. Brains were removed from the skull and post-fixed in the same fixative for 3 days shaking at 4C.

### Whole Mouse Brain Processing

After washing in 1X PBS, brains incubated in inactivation solution of 1% acrylamide (40% stock, Biorad #161-0140), 1M glycine (Sigma-Aldrich, St. Louis, MO, #G7126) in 1X PBS. After washing in 1X PBS, brains were put into clearing solution of 200mM sodium dodecyl sulfate (SDS) (Sigma-Aldrich, #L3771), 20mM lithium hydroxide monohydrate (Sigma, #254274), 40mM boric acid (Sigma-Aldrich, #7901), pH 8.5–9.0 and left shaking at 55C for 4 weeks until white matter tracts were translucent to the eye in SDS. Brains were washed in 1X PBS or Weak Binding Solution (WBS) for up to 1 week and immunolabeled with the ‘Whole Brain SWITCH for labeling intact mouse brain (Supplemental Protocol 1).

### Mouse Brain Section Processing

Where sections were analyzed, brains were sliced to 100um on a vibratome (Leica VT100S) and stored at 4C in 1X PBS + 0.02% sodium azide (Sigma-ALdrich, #S2002). For SWITCH labeling, individual sections were incubated in clearing solution shaking at 55C for 2 hours and were washed in 1X PBS. Sections were immunolabeled with the SWITCH protocol for labeling mouse brain sections (Supplemental Protocol 2).

### Human Tissue Processing

Human tissue blocks were deparaffinized by sequential immersion in xylene, ethanol, and water (details in Human SWITCH protocol). Then, blocks were incubated in 1% GA in 1X PBS for 10 days at 4C. Brains were incubated in clearing solution shaking at 55C until the tissue appeared translucent (4–8 weeks). Following clearing, tissue was labeled using the Human SWITCH protocol for labeling human autopsy specimens (Supplemental Protocol 3).

### Antibodies & Dyes

The primary antibodies used are as follows:

| Target | Host | Company | Product # | Dilution |
| --- | --- | --- | --- | --- |
| Aβ (D54D2) | Rabbit | Cell Signaling Technologies | 8243 | 1:100 –1000 |
| Myelin Basic Protein* SMI-99 | Mouse | BioLegend | 808401 | 1:50 –500 |
| Myelin Basic Protein* SMI-94 | Mouse | BioLegend | 836502 | 1:50 –500 |

\*These antibodies were used concurrently as per the manufacturer's recommendation.

#### Additional labeling reagents

Hoechst 33528 was used for nuclear labeling (Sigma #14530).

All secondary antibodies were Pre-adsorbed F(ab)2’ AlexaFluor-conjugated from AbCam.

### Whole Brain Image Acquisition

Whole brain images were acquired on a custom SPIM microscope built by H.C. During imaging, samples were illuminated with a sheet of light generated by scanning a focused beam from a light from a broad spectrum laser unit (SOLE-6 with 405, 488, 561, 647 nm, Omicron) through a low NA objective (Macro 4X/ 0.28 NA, Olympus) with a galvo-scanner (6215H, Cambridge Technology). Collection of emitted light on the microscope occurs through a long working distance high NA objective (10x/0.6 NA WD 8mm CLARITY, Olympus). The microscope is outfitted with four sCMOS cameras (Orca Flash4.0 V2, Hamamatsu) for simultaneous multichannel detection and signal recording. During acquisition, the samples were illuminated simultaneously from dual illumination arms (one on each side) to minimize the shading effects of light scattering elements in the brain. Dynamic confocal mode of detection is implemented by synchronizing the scanning speed of the illumination beam with the read-out speed of the rolling shutter mode of sCMOS camera, which improves the signal to background ratio by filtering out background noise from out-of-focal planes. Sample is mounted on a motorized stage with x, y, z translation and theta rotation (M-112.2DG, M-111K028, M-116.DG, Physik Instrumente) for mosaic imaging. Z-stack imaging by sample scanning alone is slow due to communication overhead between the host computer and the stage controller. We achieved high-speed volume imaging by scanning the light sheet along the depth direction with a galvo-scanner and synchronizing the position of the light sheet with the detection objective’s focal plane by moving the detection objective with a piezo actuator (P-628.1CL, Physik Instrumente). To maintain light sheet position on the focal plane of the objective across the entire sample volume, we implemented an image based autofocusing algorithm.^76^ The laser settings are determined such that about 5% of the images are saturated to its maximum gray level for high signal to background ratio. The sample chamber is filled with the refactive index matching solution (RIMS)^77^. Depending on the refractive index of the medium, the beam waist position of the illumination light sheet shifts along the illumination beam direction. Each illumination objective is mounted on the piezo actuator (P-628.1 CL, Physik Instrumente) to allow the beam waist position to be adjusted to the center of the detection objective. Sample chamber is specially designed to allow for free motion of the detection objective while preventing leakage of the immersion medium.

### Whole Brain Image Processing

Each tile is first corrected for the non-uniform illumination pattern using a modified algorithm from Smith and colleages^78^. Multiple stacks of acquired images are stitched with Terastitcher.^79^ Each tile has 15% overlapping area with the neighboring tiles for calculating stitching parameters. The voxel size of raw data is 0.58 x 0.58 x 5.0um. After stitching, the data set is down-sampled in X and Y dimension for further analysis with Imaris software (Bitplane).

### Section Image Acquisition

For section and human tissue imaging, tissue was mounted onto microscope slides (VWR VistaVision, VWR International, LLC, Radnor, PA) with either Fluromount G Mounting Medium (Electron Microscopy Sciences, Hatfield, PA, USA) or RIMS solution^77^.

Slice images were acquired on a Zeiss LSM Inverted 710 microscope using Zen 2012 software (Carl Zeiss Microscopy, Jena, Germany). Images with cellular resolution were taken using a C-apochromat 40X, water immersion objective, NA 1.20. Section overview images used a Plan-apochromat 5X, air objective, NA 0.16. Pinhole, optical sectioning, and laser settings were determined for each experiment and kept consistent for all images within an experiment or that were included within one graph.

### Human Brain Image Acquisition

Human brain images were acquired on a Leica TCS SP8 Confocal Microscope using LASAF software (Leica Microsystems, Wetzlar, Germany). Images were taken using a 20X 1.0NA CLARITY optimized objective with 6mm working distance. The pinhole, optical sectioning, and resolution and laser settings were empirically determined for one brain, and kept constant for imaging subsequent samples.

### 3D Image Quantification

Images were analyzed using Imaris (Bitplane, Zurich, Switzerland). All quantification steps were performed on raw images by blinded investigators. For whole brain analyses (Figure 1), each whole brain file was segmented by hand using white matter tract and regional guidelines from the Allen Brain Atlas (Allen Mouse Brain Atlas, Coronal) to delineate boundaries for each major brain region (Video 5). After segmentation, a spots object was created on a 12 month brain. The parameters were fixed over the entire brain, and spots were separated into the bounded brain regions using the Spots into Surfaces tool in Imaris Xtensions. A spots object was created on each brain and these objects were similarly split into brain regions using the Xtension. Data was exported to CSV and analyzed GraphPad Prism 7.0a for Mac. For human brain analyses (Figure 3), the 3D rendering was randomly segmented into five, 0.5mm^3^ sub-sections to account for differences in block size. An identical spots object was created within each of the five subsections, and the data was exported to CSV and analyzed GraphPad Prism 7.0a for Mac.

### 2D Image Quantification

For 2D analyses, images were imported into FIJI^80^ as TIFF files. Relevant values were exported to GraphPad for statistical analyses. For human amyloid analyses (Figure 3), two blinded observers counted plaques within images using the Multi-point tool. Data was saved as a CSV and analyzed in GraphPad Prism 7.0a for Mac. For white matter quantification (Figure 4), blinded observers analyzing only the myelin image channel of a 5X section-overview image segmented white matter tracts. Then the segmentation was overlaid on the Aβ channel as a selection, within which a threshold was applied to the images. Finally, the Analyze Particles tool was used to count individual deposits. Data was saved in a spreadsheet and analyzed in GraphPad Prism 7.0a for Mac.

### Representative Images and Videos from Whole Brain Data

Representative images from the whole-brain datasets are either 2D images of the 3D rendered whole dataset or digitally sectioned at 5–100um in Imaris using the Orthoslicer tool. Videos are created using the Key Frame Animation tool in Imaris. The brightness of the images has been individually adjusted for each brain to enhance 2D/3D viewing of specific objects. Because of these alterations, no direct comparisons of labeling intensity should be made between images.

### *In situ* hybridization probe design

RNA antisense probes were generated by PCR-amplifying human cDNA with human-specific *APP* primers with a T7 RNA polymerase recognition sequence (TAATACGACTCACTATAGGG) fused to the reverse primer (Table 1). The resulting PCR product was gel extracted and *in vitro* transcribed using a DIG-RNA labeling kit (Roche).

**Table 1.**
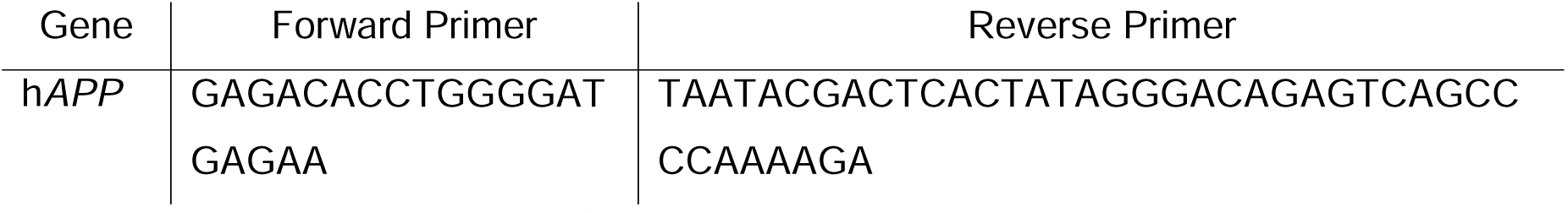
Primer sequences (5’–3’) for in situ probe preparation.

### Immuno-in *situ* hybridization (Immuno-ISH)

Mice were anesthetized by isoflurane in an open system and perfused with RNase-free PBS followed by RNase-free 4% formaldehyde. Brains were dissected, drop fixed in RNase-free 4% formaldehyde for 12 hours, equilibrated in 30% sucrose-PBS, and frozen in O.C.T. (TissueTek). Cryosections (10μm) were incubated with a DIG-labeled RNA antisense probe (1:1000 in hybridization buffer) overnight at 65^o^C, washed in 1X SSC/50% formamide/0.1% Tween-20 3X 30 minutes at 65^o^C followed by 1X MABT for 30 minutes at room temperature. Sections were blocked with 20% heat-inactivated sheep serum/2% blocking reagent (Roche)/1X MABT for 1 hour and then incubated with mouse anti-DIG antibody (Roche; 1:2000) and rabbit anti-amyloid β (Cell Signaling; 1:500) diluted in blocking solution overnight. Sections were washed with 1XMABT 2X 20 minutes, incubated with donkey anti-rabbit Alexa-488 (Invitrogen; 1:1000) diluted in blocking solution for 1 hour, and washed with 1XMABT 5X 20 minutes. Sections were then prestained with 100mM NaCl/50mM MgCl_2_/100mM Tris pH 9.5/0.1% Tween-20 2X 10 minutes, followed by staining with NBT/BCIP (Roche; 4.5 μl/ml and 3.5 μl/ml, respectively, in prestaining buffer) for 2 hours. Sections were washed with 1X PBS 3X 15 minutes, incubated in xylene 3X 5 minutes, and mounted with VectaMount (Vector Laboratories).

### Ex vivo electrophysiology

Acute hippocampal slices were prepared from C57bl/6 mice aged2-2.5 months old which are wild type group and 5xFAD group. The mice were anesthetized with isoflurane and decapitated. The experimenter was blinded to the group of animal. Transverse brain slices (250 μm thick) were prepared in ice-cold dissection buffer (in mM: 211 sucrose, 3.3 KCl, 1.3 NaH2PO4, 0.5 CaCl2, 10 MgCl2, 26 NaHCO3 and 11 glucose) using a Leica VT1000S vibratome (Leica). Slices were recovered in a submerged chamber with 95% O2/5% CO2-saturated artificial cerebrospinal fluid (ACSF) consisting of (in mM) 124 NaCl, 3.3 KCl, 1.3 NaH2PO4, 2.5 CaCl2, 1.5 MgCl2, 26 NaHCO3 and 11 glucose for 1 h at 28–30 °C. Whole cell patch clamp with current steps from 0 pA to +130 pA at 10 pA increments for 800ms were used to verify the ability to form action potential (AP) on HEKA ECP 10 Amplifier (Germany) with Pulse software. Number of APs at each current step; Resting membrane potential (RMP); Threshold of APs; Amplitude of APs from RMP and threshould as well as fast afterhyperpolarization(fAHP) were measured and analyzed.

### Statistics

All statistics were performed in MatLab or GraphPad Prism. Individual statistical tests are indicated in the text and/or figure legends for the appropriate experiments. Graphs were created in the respective analytical software packages and exported as .TIFF, .PDF, or .JPEG for inclusion in the document.

## Acknowledgements

The authors would like to acknowledge Michiel Kooreman at the Netherlands Brain Bank for help procuring the PPFE human tissue and Teresa Lima for her expert advice in preparing the human PPFE brain tissue for the optimized CLARITY protocol. We would also like to thank Nina Dedic for her advice and helpful comments on the project and manuscript. Additionally, we would like to acknowledge Naveed Bakh, Sung-Yon Kim, and Kamilla Tekiela for helping lay the groundwork for the experiments presented in this work.

## Funding

This work was funded by NIH grant RF1AG047661 to L.-H.T. and by the Norman B. Leventhal and Barbara Weedon fellowships to R.G.C. Additionally, K.C. was supported by Burroughs Wellcome Fund Career Awards at the Scientific Interface, the Searle Scholars Program, Packard award in Science and Engineering, NARSAD Young Investigator Award, JPB Foundation (PIIF and PNDRF), NCSOFT Cultural Foundation, and NIH (1-U01-NS090473-01).

Resources that may help enable general users to establish the methodology are freely available online (http://www.chunglabresources.org).

---

**Supplemental Figure 1.**
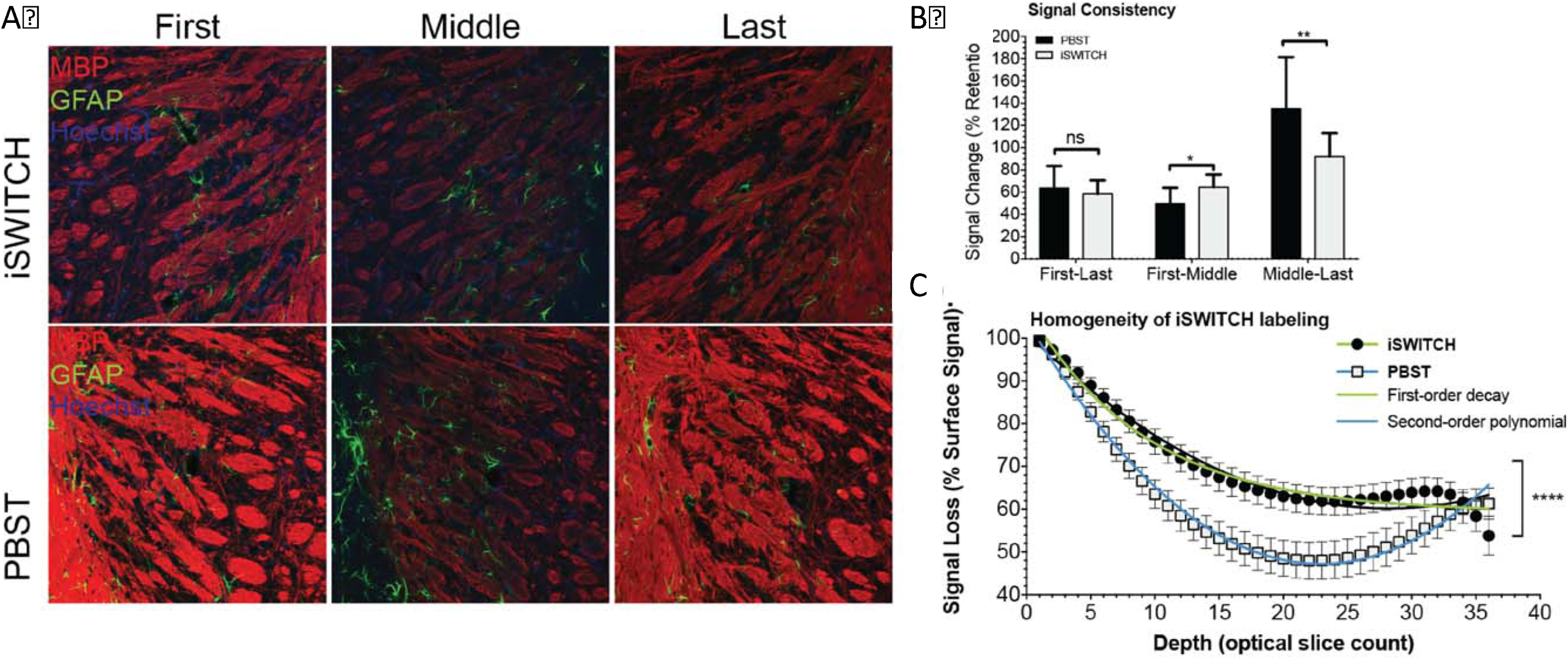
Optimized iSWITCH enhances immunolabeling homogeneity in 100um sections. A) Representative optical sections from the first (left), middle (center), and last (right) slices of z-stack image files taken from tissue labeled for myelin basic protein (MBP, red) glial fibrillary acidic protein (GFAP, green), and with Hoechst dye (blue) using either iSWITCH buffers (top) or tradititional labeling buffer phosphate buffered saline with triton x-100 (PBST, bottom). B) Quantification of MBP labeling demonstrating signal inensity consistency between the first and last sections under both conditions (left), but enhanced middle section immunolabeling signal when compared to either the intensity of the first (center) or last (right) section. C) Immunofluorescent signal loss throughout the depth of the sections is first order decay using iSWITCH buffers, indicating relatively homogenous signal with intensity that decays with depth due to laser attenuation, but using PBST the loss is a second order polynomial, indicating non-homogenous signal intensity through the thickness of the tissue. *p ≤ 0.05; **p ≤ 0.01; ****p≤ 0.0001; Graphs report Mean ± Standard Deviation.

**Supplemental Figure 2.**
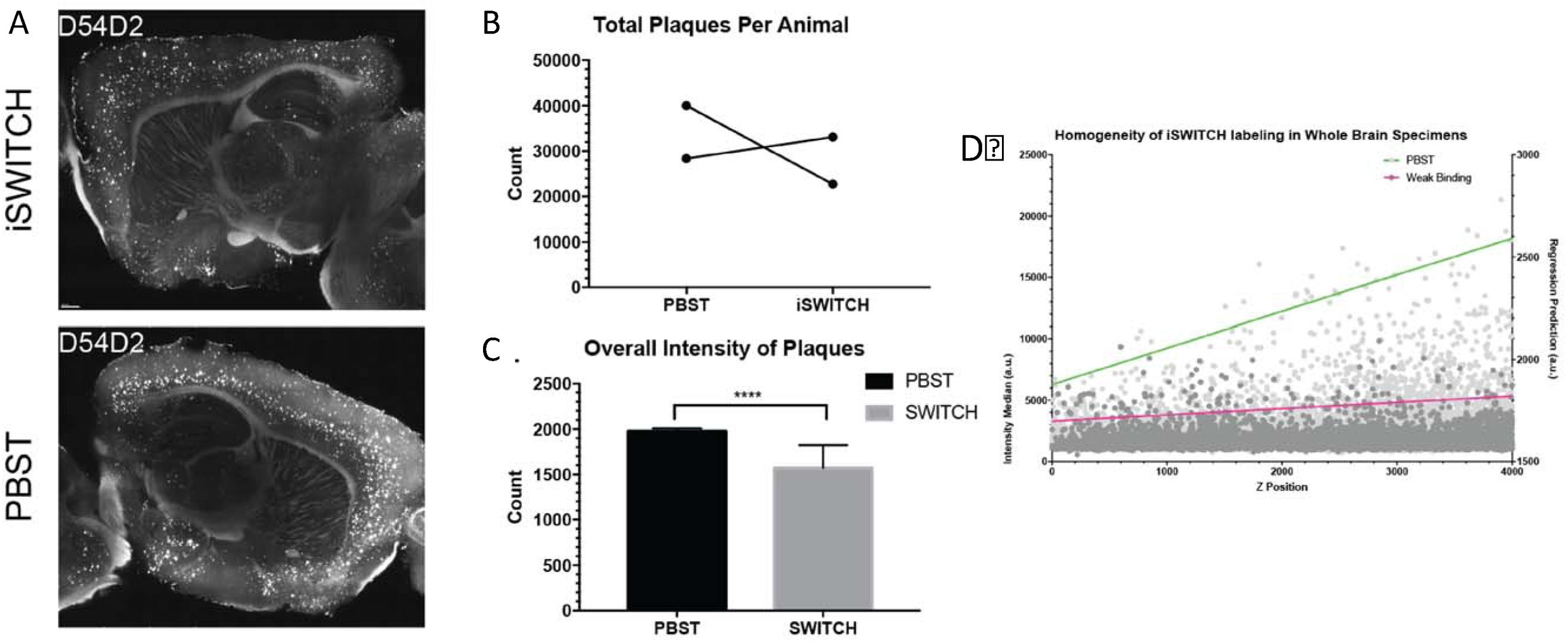
iSWITCH buffer immunolabeling whole brain for β-amyloid achieves homogenous signal. Aged 5XFAD brain was split into two hemispheres and labeled with either iSWITCH buffers or PBST. A) Representative images from the center section of each hemisphere. B) Demonstration of the consistency of overall aggregate counts between animals and no consistent effect of PBST or iSWITCH on aggregate detection. C) Aggregates labeled by iSWITCH show overall lower intensity, which may be driven by more antibody availability for surface antigens. D) Plot of the intensity of aggregates detected through sample (left axis) with regression (right axis) showing non-homogenous signal intensities in PBST labeled brains, which is rectified using iSWITCH buffers. Graphs report Mean ± Standard Deviation unless otherwise noted in the text.

**Supplemental Figure 3.**
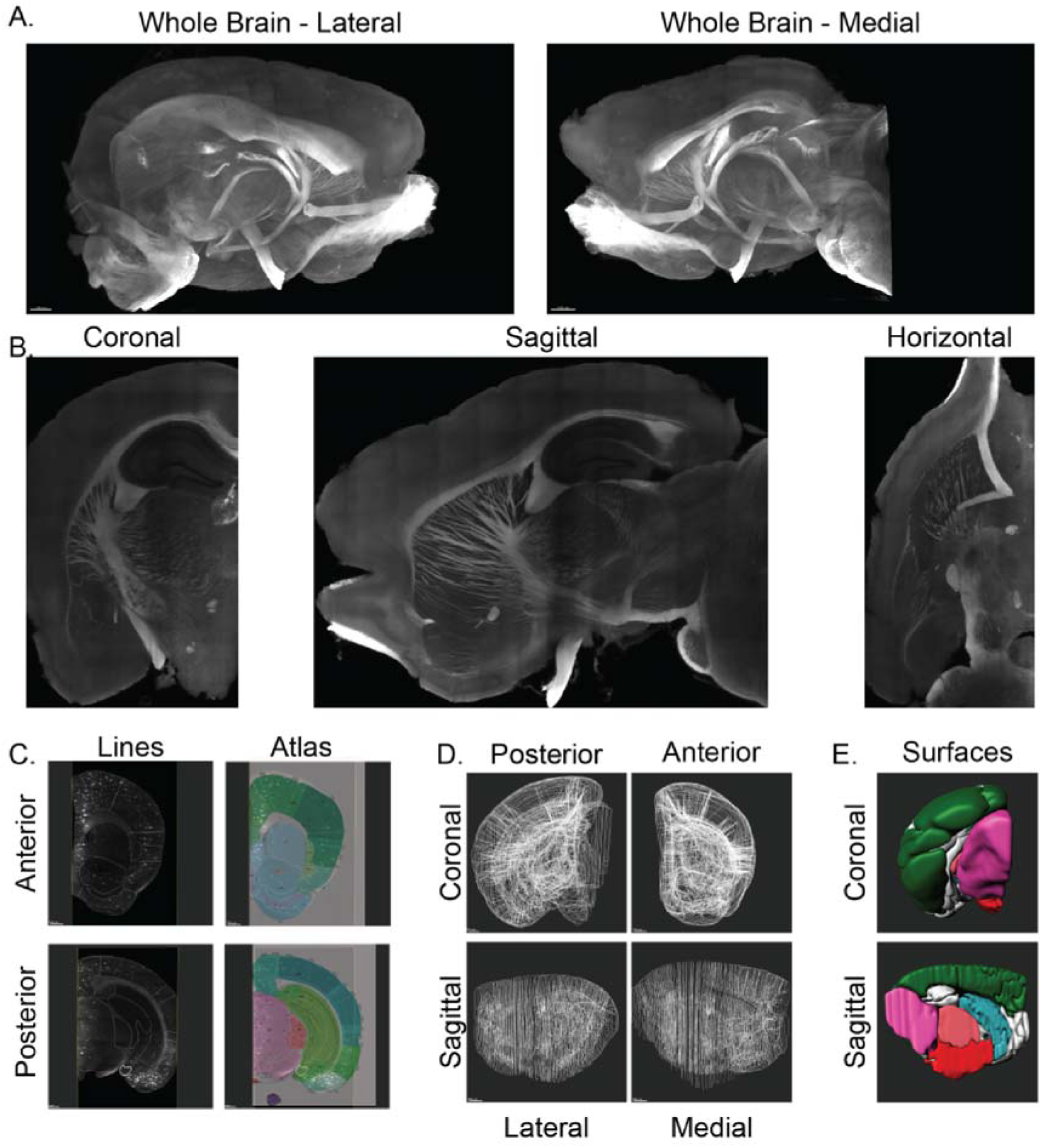
White-matter based hand segmentation of brain areas provides robust identification of specific regions. A) 3D rendering of white matter tracts in the whole mouse brain. B) Optical slice-wise visualization of white matter tracts. C) Representative examples of hand-segmentation (left) next to the Allen Mouse Brain Atlas section (right, anterior section #43, posterior section #85) from which they were drawn. Website: © 2015 Allen Institute for Brain Science. Allen Mouse Brain Atlas [Internet], Available from: http://mouse.brain-map.org^311^ D) Representative images showing the whole-brain segmentation. E) Representative images of the volume-rendered segmentation.

**Supplemental Figure 4.**
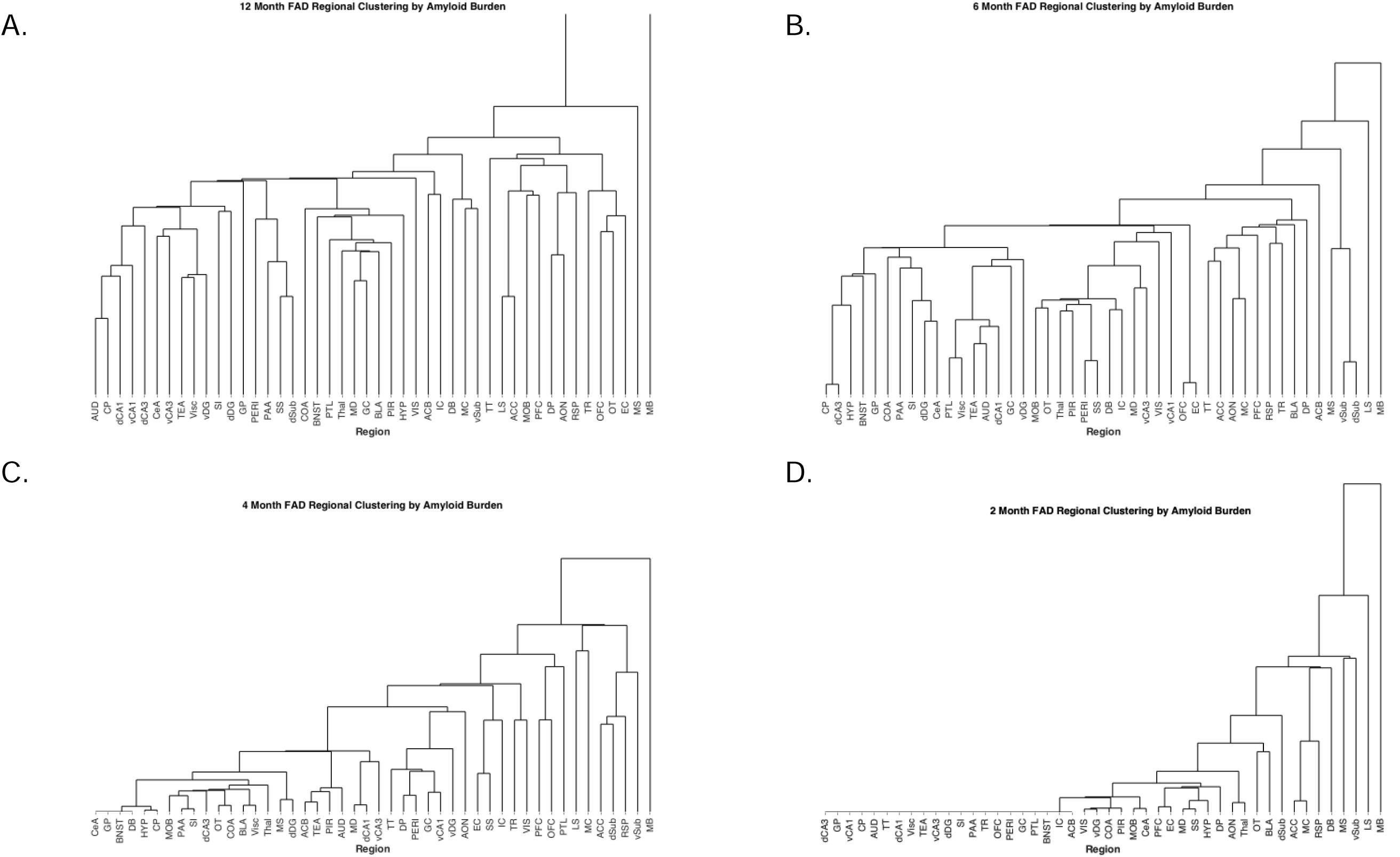
Imaging reveals no correlation between transgenic mRNA expression and amyloid aggregates. A) Representative images of the double *in situ* hybridization and immunofluorescence. B) Brain-wide correlation graph demonstration lack of correlation between signals. C) Correlation coefficients split by region and displayed with p-value.

**Supplemental Figure 5.**
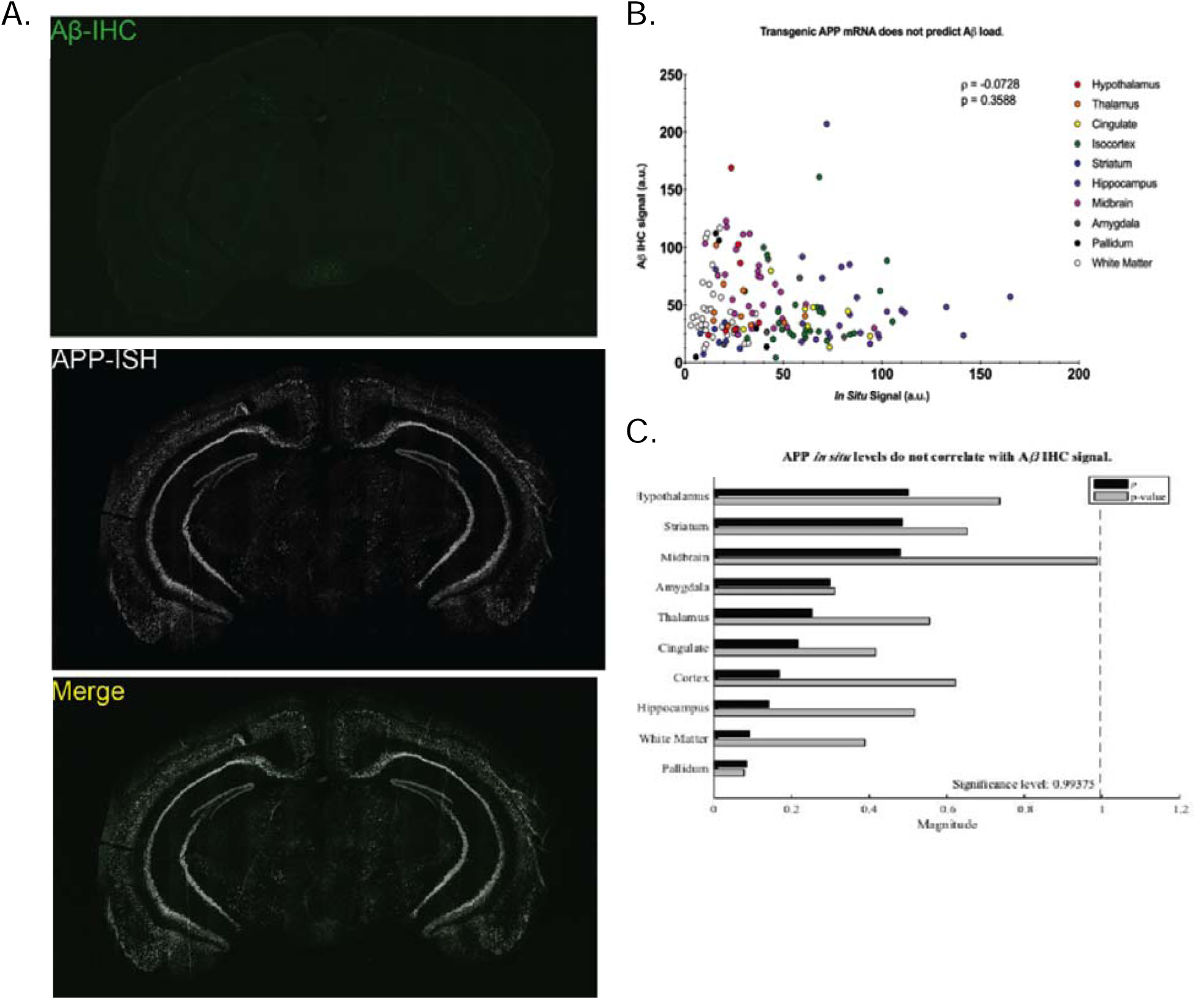
Hierarchical clustering at each age reveals amyloid deposition within networks appears with age. A-D) Dendrograms of hierarchical clustering of amyloid density at each age examined in the 5XFAD mice. A) 12 months. B) 6 months C) 4 months and D) 2 months aged animals.

